# Active neural coordination of motor behaviors with internal states

**DOI:** 10.1101/2021.12.10.472142

**Authors:** Yisi S. Zhang, Daniel Y. Takahashi, Ahmed El Hady, Diana A. Liao, Asif A. Ghazanfar

## Abstract

The brain continuously coordinates skeletomuscular movements with internal physiological states like arousal, but how is this coordination achieved? One possibility is that brain simply reacts to changes in external and/or internal signals. Another possibility is that it is actively coordinating both external and internal activities. We used functional ultrasound imaging to capture a large medial section of the brain, including multiple cortical and subcortical areas, in marmoset monkeys while monitoring their spontaneous movements and cardiac activity. By analyzing the causal ordering of these different time-series, we found that information flowing from the brain to movements and heart rate fluctuations were significantly greater than in the opposite direction. The brain areas involved in this external versus internal coordination were spatially distinct but also extensively interconnected. Temporally, the brain alternated between network states for this regulation. These findings suggest that the brain’s dynamics actively and efficiently coordinate motor behavior with internal physiology.

---

Animals and humans continuously regulate physiology and behavior to maintain stability, i.e., keep physiological variables within a tenable range (1). This regulation not only involves triggering autonomic reflexes that directly adjust physiological processes such as heart rate, glucose level, and body temperature but also directs skeletomotor behaviors to interact with the external world that affect physiological states (e.g., feeding, locomotion, social interaction)(2, 3). The changing internal states of the body remodel sensorimotor interactions with the external environment on various timescales (4). For example, the phase of the cardiac cycle influences the emotional processing of faces (5), vocal interactions are correlated with autonomic oscillations (6–8), and hunger can modulate the switch between sleep versus foraging behaviors (9). In all cases, the internal physiological states must be coordinated with motor behaviors through the dynamics of large-scale networks of cortical and subcortical regions, but how?

Based on the spectrum of cytoarchitectonic differentiation, one proposal is that the mammalian brain follows a ‘centrifugal’ cortical organization from the outer side (the primary sensory and motor areas) to the inside. Inside areas include the heteromodal, paralimbic and limbic regions that, in humans, overlap substantially with the default mode network (10–12). The outside areas directly regulate the interactions with the external environment, while the inside areas are associated with autonomic functions regulating the internal milieu through subcortical areas, primarily the hypothalamus (12–14). A more recent proposal, based on the meta-analyses of rat cortical neuroanatomy, is a topologic core-shell arrangement with two sensory-motor modules (core) and two limbic modules (shell) corresponding to this external-internal dichotomy (15). In humans, functional imaging and gene expression analyses also reveal a division of the brain into two cortical networks (16, 17). Such a brain architecture entails an embodied account of perception, emotion, and decision making (18–23).

However, within such a framework, how are motor behaviors and the internal state of the body organized with the brain’s activity in space and time? If we assume a context in which there are no overt external sensory signals, and the animal coordinates interactions with external and internal environments by its intrinsic dynamics, then there are currently three hypotheses for the inter-relationships between brain activity, motor behavior, and internal physiology (Fig. 1A). The first hypothesis postulates that spontaneous motor behaviors--like twitches, facial movements, and fidgets--drive the large-scale activity of the brain (24–27). The behavior-related wide-spread brain activity could originate from re-afferent sensory input and may facilitate contextualized signal processing (28). Under this hypothesis, there are two possibilities for the internal physiological state: it is either a follower of movements through peripheral regulation (Hypothesis #1.1) or centrally regulated (Hypothesis #1.2). In the second hypothesis, brain dynamics are passive responses to ongoing physiological states and provide signals--including those to produce behaviors--to maintain homeostasis of the body (Hypothesis #2) (29–31). The third hypothesis proposes that the brain not only actively predicts signals of the external world (32, 33) but also the interoceptive signals of the body for homeostatic control (Hypothesis #3) (34–36).

**Figure 1.**
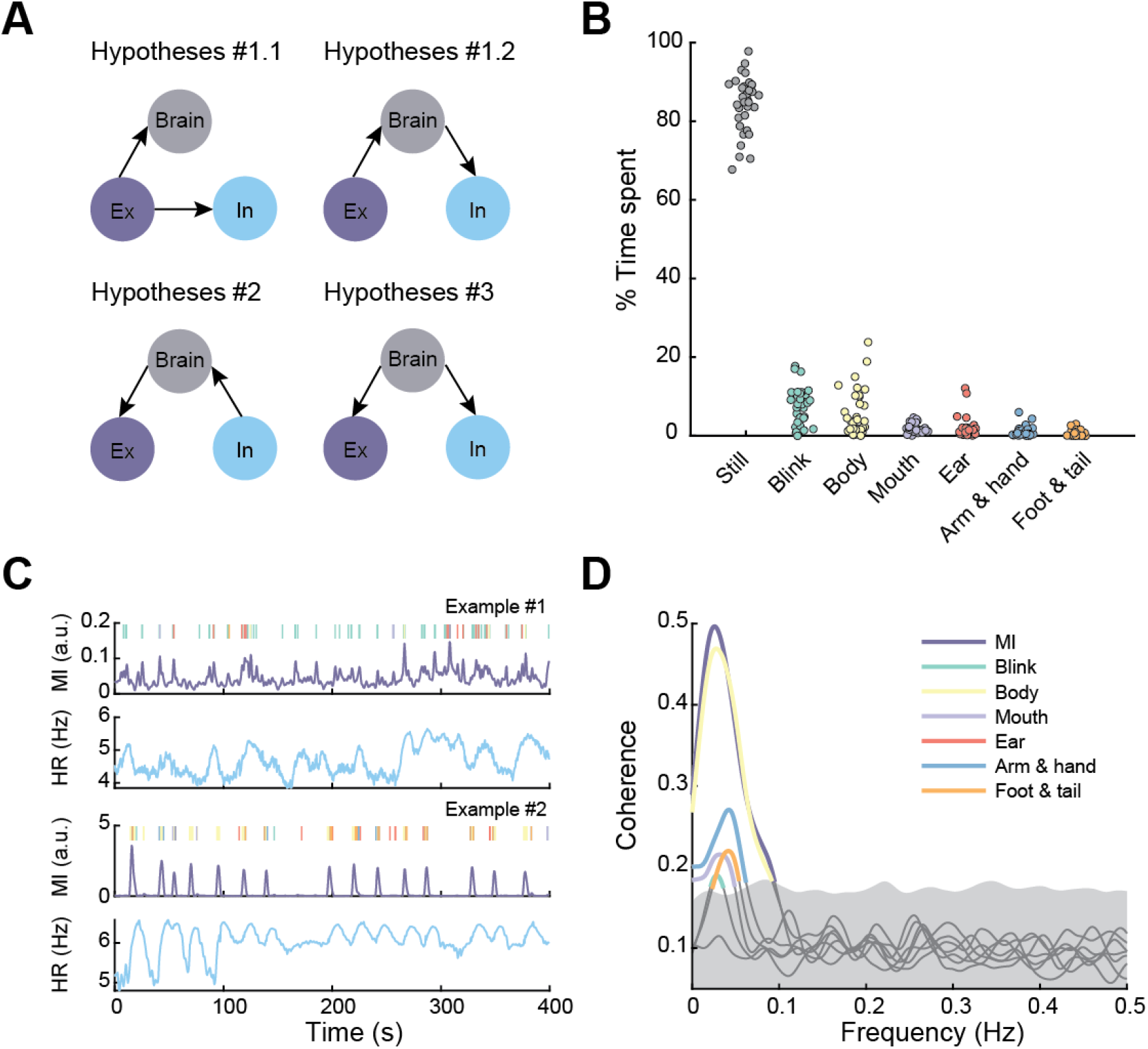
Organization of external and internal activities. (A) Candidate hypotheses on the interrelationships between brain activity, movement and heart rate fluctuation. (B) Percentage time spent on different types of movements. Each point is a session. Jitters are added to help visualize. (C) Two exemplar data of movement type, MI, and HR. Types of movements are color-coded similarly to (B). Note that small MI variations correspond to facial movements such as blinks and large MIs correspond to movements of the body parts. (D) Coherence between each HR-movement type pair and the HR-MI pair. Highlighted segments are significantly higher than the 95% CI of the phase-randomized surrogates.

To test these hypotheses, we studied the inter-relationships between the brain, external (spontaneous movement), and internal (cardiac) activities in marmoset monkeys (*Callithrix jacchus*) under a task-absent context. These various activities are prevalent on the timescale of tens of seconds (24, 37–41), and thus can be tested in experimental sessions lasting for 10-20 minutes. We simultaneously used functional ultrasound (fUS) imaging of the midsagittal plane of the marmoset brain, along with videos of behaviors and measures of heart rate (electrocardiogram, ECG) to establish: 1) the direction of information flow between the three components, and 2) the spatial and the temporal organizations of brain activity as a function of spontaneous movements and cardiac changes.

## Results

Subjects (n=3) were placed in a partial restraint device to allow for both stable neural imaging and movements of most of the body. Using behavioral videography analysis (35 sessions, 27259 seconds of recording), we detected occasional movements of the limbs, tail, and body, as well as facial movements, including blinks and movements of ear and mouth (Fig. 1B). Movements were also quantified by the motion intensity (MI) regardless of the type. In parallel, we carried out electrocardiography (ECG) for continuous heart rate recording (Fig. 1C). The occurrence of different behaviors varied coherently with heart rate to different degrees with all, except ear movements, showing a peak around 0.03 Hz (test against the 95% confidence interval (CI) of phase-randomized surrogates; Fig. 1D). Using MI alone, the coherence between movement and heart rate could also be well captured (Fig. 1D), and there was no temporal difference between MI and elaborated movements with respect to heart rate (Fig. S1). Thus, for ease of presentation, we used MI to represent all kinds of movements in the following analyses.

Meanwhile, we measured brain activity in a large portion of the midsagittal plane, including cortical and subcortical areas using fUS (35 sessions; Fig. 2A). fUS measures cerebral blood volume (CBV) dynamics in the microvasculature, an indirect measure of neuronal activity(42–44), with a large field of view (FOV) (16 mm (AP) × 20 mm (DV)), a spatial resolution of 130 μm (AP) × 125 μm (DV), and a frame rate of 2 Hz. The FOV covered primary sensory and motor areas: the medial motor (M1) and somatosensory (SS); motor associate areas: pre-supplementary motor area (preSMA) and supplemental motor area (SMA); high-heteromodal and paralimbic areas: prefrontal (PFC), cingulate motor area (CMA); as well as cortices constituting the DMN: the ventromedial prefrontal cortex (vmPFC) and posterior cingulate cortex (PCC) (45). It also covered subcortical areas relevant to autonomic functions(46), including the dorsal and ventral lateral septal nuclei (LSD and LSV), preoptic area (POA), ventromedial hypothalamus (VMH), posterior hypothalamus (PH), mediodorsal nucleus of the thalamus (MD), habenula nuclei (Hb) and part of the reticular formation (RF) (Fig. 2A). Thus, the simultaneous activities of many brain areas (though not all of them) highly relevant to motor behavior and internal physiology were incorporated in the measurement.

**Figure 2.**
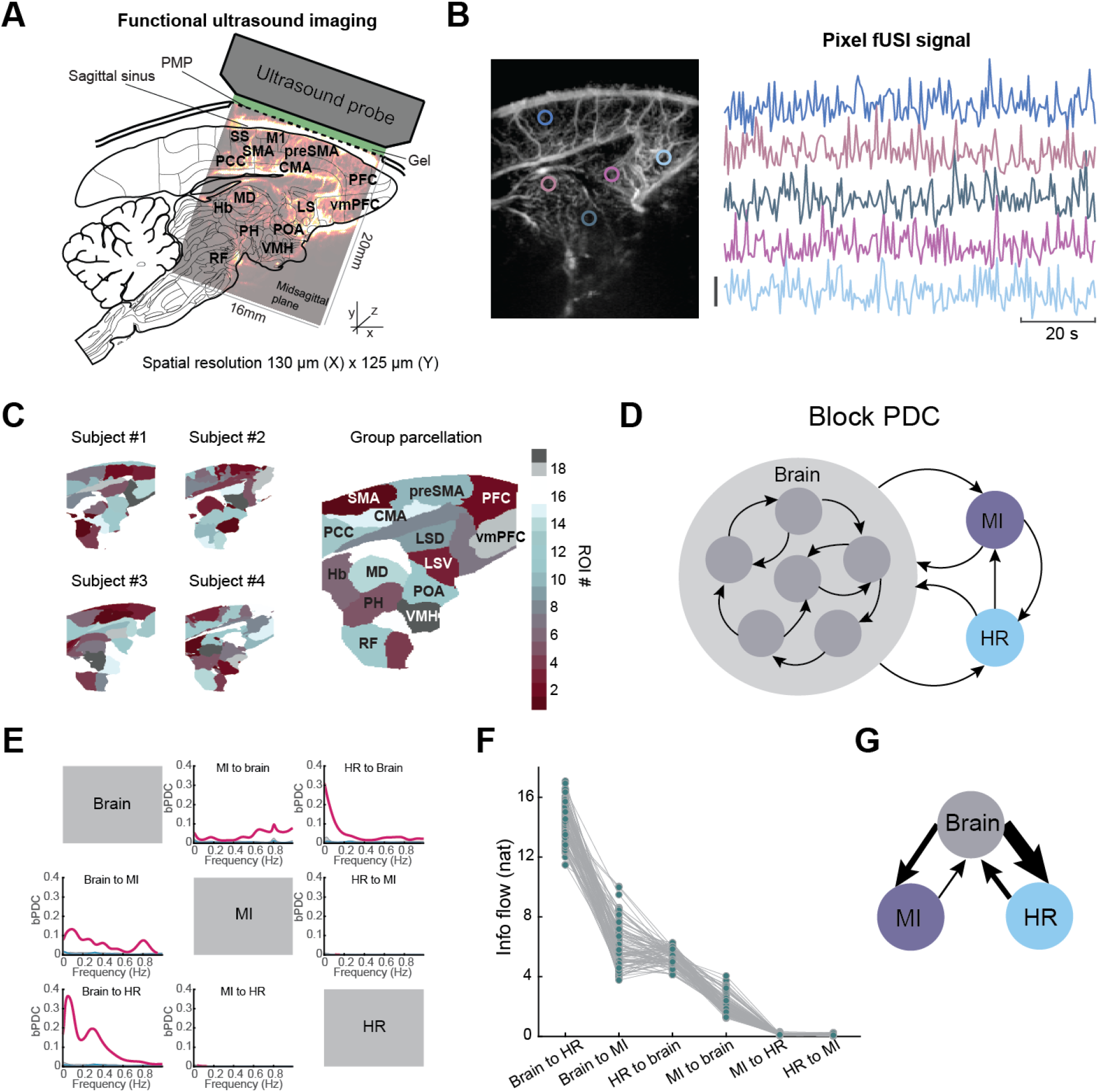
The brain plays an active role in coordinating movement and heart rate. (A) fUS setup and brain regions covered by imaging. (B) Exemplar pixel signals from different areas of the brain. Vertical bar indicates 2 standard deviation. (C) Within subject and group-level brain parcellation. We use SMA to represent the ROI covering SS, M1, and SMA. LSD: lateral septal nucleus dorsal; LSV: lateral septal nucleus ventral. (D) Illustration of bPDC. (E) bPDC of each direction. Red means significantly higher than phase-randomized surrogate distribution. Note that the lines in “HR to MI” and “MI to HR” cells are very low. (F) Estimated distribution of information flow of each direction, sorted from high to low. Outward flow from brain is greater than the corresponding inward flow (p<0.05 Bonferroni corrected). (G) Information flow graph. The thickness of the arrows is proportional to information flow.

To evaluate the relationship between brain and peripheral activities, we first parcellated the image into regions of interest (ROIs) based on fUS signals. We estimated a group-level parcellation using all available fUS data by concatenating temporally reduced data using principal component analysis (4 subjects, 60 sessions, 85427 frames; Methods), and segregating the brain section into 19 ROIs by spectral clustering (Fig. 2B, C). Among the 19 ROIs, 14 were identified as non-vessel ROIs and were named based on corresponding anatomical annotations after aligning to a reference brain atlas (Methods; Fig. 2C, Fig. S2) (47, 48). The brain parcellation results were consistent across subjects (Fig. 2C), and thus we concatenated the ROI signals across all sessions.

The information flow between time-series of brain activity, movements (MI), and heart rate was assessed using block partial directed coherence (bPDC), a measure of information flow in the frequency domain (49) (Methods) and invariant to causal filtering like the hemodynamic response function (see Supplementary text) (50). The bPDC treated the ROI signals as one multivariate time-series network node and the movement and heart rate as the other two nodes (Fig. 2D). We found that the bPDC was significant in both directions between brain and movement, and between brain and heart rate; while the bPDC between movement and heart rate was not significant, suggesting that their coordination was via the brain (test against the 95%confidence intervals (CI) of phase-randomized surrogates, Bonferroni corrected; Fig. 2E). Hence, we could reject Hypothesis #1.1 as the information flow between movement and heart rate was indirect. We then tested if the brain activity was driven more by the periphery or vice versa by comparing the total information flow in each direction, which was calculated by integrating the bPDC over frequencies (Methods). On average, information flowing from the brain to the other two components was significantly higher than the inward flow (bootstrap and paired z-test, p<0.05 Bonferroni corrected; Fig. 2F,G; regardless of whether filters were applied (Fig. S3)). Consistently, brain activity led the other two activities temporally, and the causal direction was due to 0 – 1 s leading of the brain (Fig. S4). The higher outward information flow was observed for all types of movements as well (Fig. S5). Thus, the results favored Hypothesis #3, that the brain activity was mostly predictive of motor output and fluctuations of internal physiology.

A caveat is that brain hemodynamics is naturally linked to cardiac activity, and one may expect to observe information flow solely from heart rate to brain (51). While the influence from heart rate to CBV could be added up by the effect of blood flow, we argue here that information flow in the opposite direction can be a conservative estimation of neural regulation of heart rate. First, for reality check, we randomly selected patches within the superior sagittal sinus, an area outside the neural content as a control, and assessed the information flow within a brain-sinus-HR network. The information was significantly greater flowing from heart rate to sinus than the opposite direction (n=30, p=0.0036, paired t-test), verifying that blood flow not driven by neural activity was influenced by heart rate. The information flow from CBV to heart rate can be realized via hidden neural activity in the neighborhood of the CBV signal. Consider a network consisting of the CBV of an ROI, the neuronal activity of this ROI, and heart rate, where the neural activity causes an immediate CBV change via neurovascular coupling as well as a heart rate change with a relatively longer latency via signaling pathway to the heart. Meanwhile, the heart rate influences brain hemodynamics (Fig. S6A). If we only measure the CBV and heart rate, it is possible to observe information flow from CBV to heart rate as long as the weight from neuron to CBV is greater than that from heart rate (Fig. S6B). Thus, using CBV as a surrogate for neural activity, it is possible to reveal the neural influence on heart rate, and our results should indicate this.

One possible mechanism coupling external and internal activities is that they could be driven by a common set of brain regions; alternatively, they could be influenced by separate regions that are themselves temporally coordinated (Fig. 3A). To test this, we first established the network composed of individual ROIs, movement, and heart rate using PDC analysis for univariate time-series (Fig. 3B) (52). The strength of connection in each direction between two nodes was quantified as integrated information flow. Two different subsets of ROIs were involved in strong information exchange with movement and heart rate (Fig. 3C, D). For example, the premotor areas SMA and CMA are known for motor sequence and planning (53, 54); the PDC analysis revealed their information exchange with movement. Another area exhibiting highly predictive activity of movement was the RF, whose function of coordinating crucial somatic movements as well as relaying descending signals to motor neurons has been well documented (54). The areas that sent significant information to heart rate including the LSD and VMH are also known to regulate motivation in general and cardiovascular functions (55). Furthermore, the estimated information flow between brain and heart rate was not driven by different regional density of vascularization; in both directions, the information flow was not dependent on the vascular density (p>0.05; Fig. S7). To further verify our results based on functional parcellation, we aligned the fUS image to a reference atlas (Methods) (48) and conducted PDC analysis with anatomical segmentation. The significant information flow from SMA, CMA, and RF to movement (MI), and from LSD and VMH to heart rate (HR) were robust against parcellation method (Fig. S8). Thus, movement and heart rate activities were coordinated by spatially separate brain regions, with the exceptions of PH and SMA that coordinated both types of activities.

**Figure 3.**
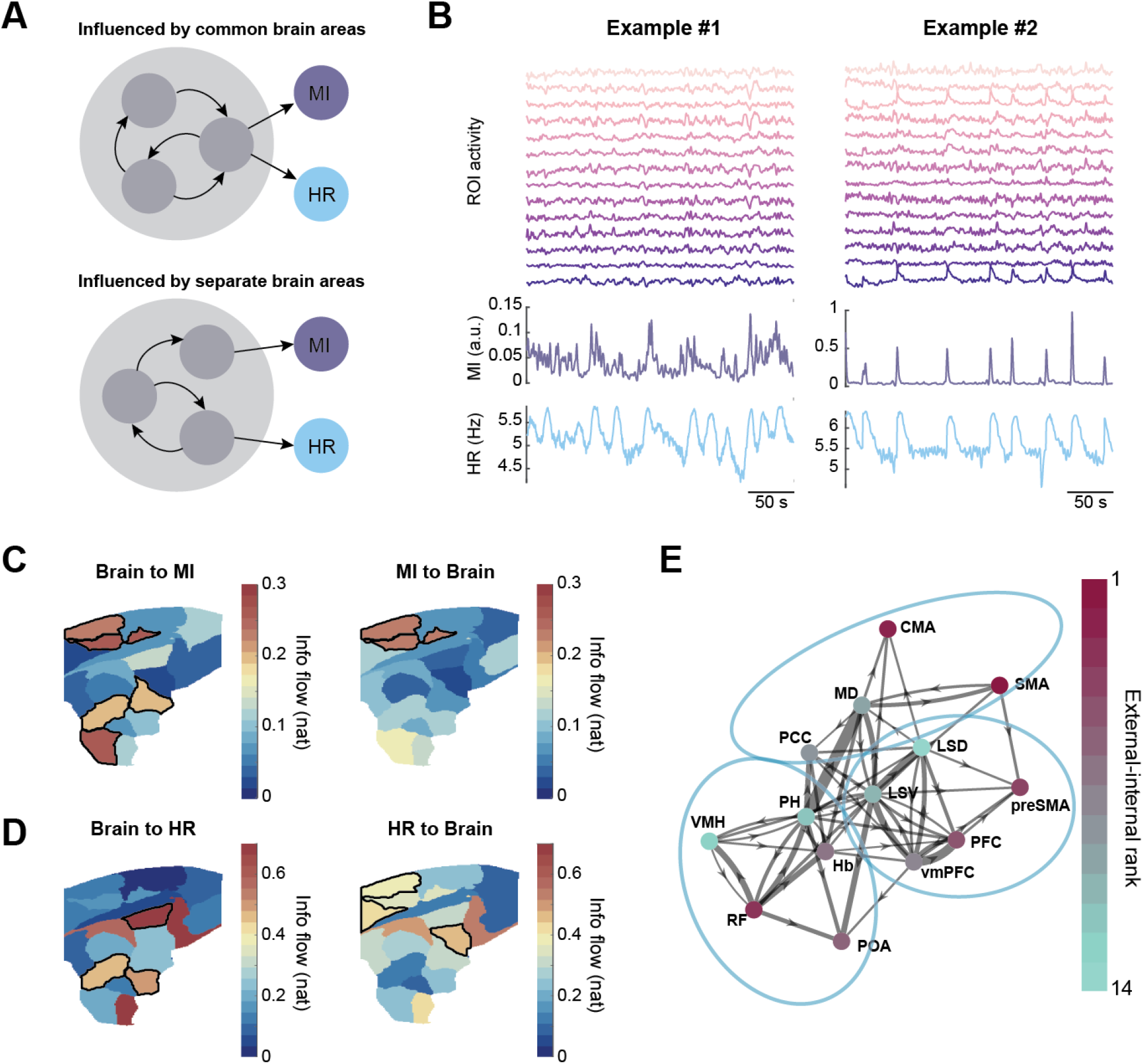
Separate brain regions regulate internal and external activities. (A) Hypotheses of spatial distribution for different brain areas involved in movement and heart rate regulation. (B) Exemplars showing individual ROI signals with MI and HR. Each time series is used as a node for PDC analysis. (C, D) Brain regions sending or receiving information flow to MI and HR. Outlined areas are the ones above the 75th-percentile of the information flow within the category. (E) Brain network summary. Circled areas are communities determined by spectral clustering. Node color corresponds to the preference of control with purple towards external activity and green towards internal activity. Note that low-level subcortical nodes are separated from higher level nodes, which are further separated into two groups, but opposite control preferences can exist in the same community.

We next investigated the connectivity between these two wide-spread groups of regions. One hypothesis is that the functional connectivity, defined as information flow between ROIs, exhibits two network communities corresponding to the external and internal control. Conversely, these two groups of regions might be so coupled that the network cannot be cleanly separated by the degree of participation in external and internal control. As the outward information flow was dominant (and also because the heart to brain direction was confounded by vascular information flow), we formulated the ROI function as the difference between information flowing to movement and to heart rate (termed control preference). The graph revealed that ROIs with different control preference were highly interconnected and could not be separated by their functional connectivity (spectral clustering for directed graph; Fig. 3E). This suggested that the coordination between movement and heart rate was due to strongly coupled brain areas. The functional connectivity established using fUS was backed by structural connectivity from fiber tractography mapped to the same areas (functional and structural connectivity correlation r=0.46, p=5.0×10^-6^; Fig. S9), comparable to measurements in other modalities and species (56–58). This internally connected network ranging from the primary sensorimotor cortex for external control to the limbic areas (here mainly the LS) for internal control is consistent with the idea of ‘centrifugal’ brain organization (12).

We then investigated how the control of ongoing behavior and internal physiology were coordinated temporally. Brain regions collectively transition into different states (39, 59), so the regulation of external and internal activities may occur concurrently during certain states or alternatively between states. In the first case, brain regions activated during the same brain state can have different control preferences, while in the second case, areas with different control preferences are correlated with different brain states (Fig. 4A). To test these possibilities, we evaluated the fluctuation of brain states by first transforming the fUS images into a spatial independent component (ICA; of *N_IC_* dimensions) – wavelet (of *N_f_* dimensions) representation (Fig. 4B). Briefly, each frame was reduced to *N_IC_N_f_* = 400 dimensions containing spatial and spectral information of the fUS signal (60). Before identifying brain states, we conducted data harmonization to mitigate the session-dependent variance such that the subsequent clustering would be driven by brain activity patterns instead of experiments (Methods; Fig. S10). Using k-means clustering with each timepoint as a sample, we clustered the reduced data into four states (n=85427 frames; Fig. 4C). The four clusters represented distinct spectral and spatial activation patterns (Fig. 4D). The fUS signals were then converted to sequences of brain states (Fig. 4E).

**Figure 4.**
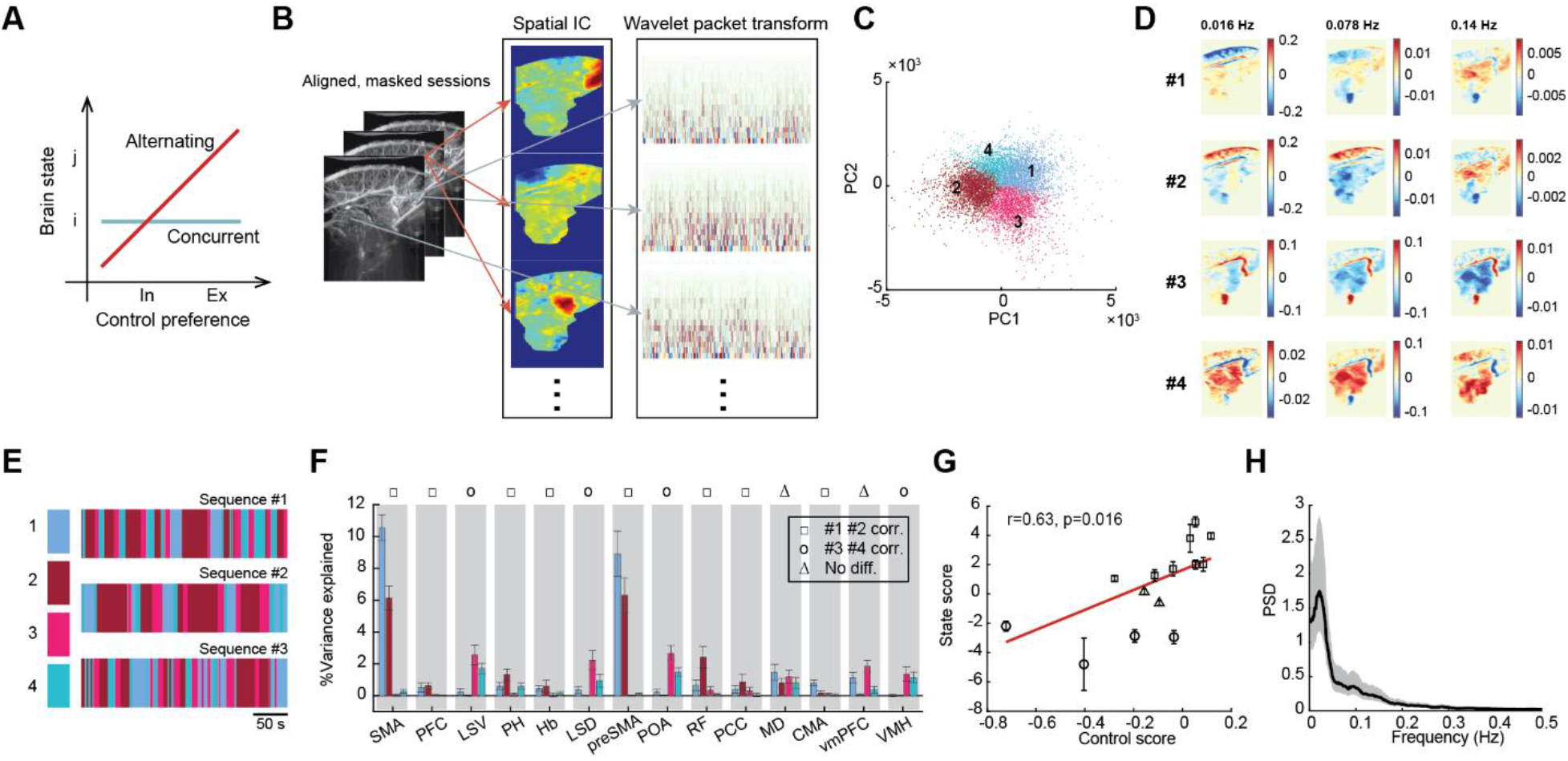
Brain alternates the control for external and internal activities. (A) Two possibilities for external and internal regulation relative to brain states: concurrent or alternating. (B) Illustration of dimensionality reduction onto ICA-wavelet space. (C) K-means clustering of the reduced data into four clusters, plotted in the space of the first two PCs for visualization. (D) Maps of mean wavelet amplitude for each cluster at different frequency bands. (E) Exemplar sequences of brain states. (F) Percentage variance explained for each ROI by different brain states. Most ROIs are correlated with either #1 & #2 or #3 & #4. Error bars are standard deviations. (G) Correlation between state score and control preference. Error bars are standard deviations. The line is fitted by weighted linear regression. r is weight correlation coefficient, and p is the p-value of the slope. (H) Power spectral density (PSD) of the alternation between #1 plus #2 and #3 plus #4. Shaded area is 95% CI. Peak around 0.02 Hz.

We then evaluated the ROIs’ participation in brain states by calculating the percentage variance of an ROI signal explained solely by the occurrence of a brain state. Most of the ROIs were significantly explained by either only states #1 and #2 or only states #3 and #4 except the MD and vmPFC (z-test, p<0.05; Fig. 4F). Based on this characteristic, we quantified a state score as the log-ratio between the total variance explained by #1 and #2, and by #3 and #4. The state score correlated significantly with control preference (r=0.63, p<0.05; Fig. 4G), suggesting that external and internal regulation occurred during different brain states with #1 and #2 for external regulation, and #3 and #4 for internal regulation. To verify this result, we also carried out spatial-spectral decomposition of fUS images using an ICA – multitaper approach (61) and obtained a significant correlation between state score and control score, demonstrating that the finding was not method specific (Fig. S11). By calculating the power spectral density of the alternation between external (#1 plus #2) and internal (#3 plus #4) brain states, we observed a peak around 0.02 Hz (Fig. 4H), close to the frequency of the peak coherence between movement and heart rate (Fig. 1D). Therefore, the coordinated external and internal activities were due to the slow oscillations between these two brain states.

## Discussion

Our findings that brain dynamics are predictive of motor activities and heart rate fluctuations support a view of the brain as being actively coordinating behavior with internal states (2, 3, 34). The regulation of these two aspects is linked to spatially separate but functionally and structurally interconnected brain regions. This organization of behavior and internal physiology with brain dynamics is analogous to the blood circulatory system: Different chambers of the heart are involved in the systemic (analogous to the external interaction) and pulmonary (analogous to the internal interaction) circulations but are also synchronized by the muscle contractions of the heart. Failure of any loop will lead to the breakdown of the entire system. Although the brain is a much more complex organ, it may drive coupled interactions with the external and internal environments in a similar way. We also found that the transitions of brain states signal the switches between external and internal control. As the brain is metabolically expensive, multifunctionality in a task-free context would be less economical (62, 63); functional alternation might be a solution for lowering the running cost of the brain (64).

Our findings suggest that in the low-frequency band (<0.5 Hz), information flow from brain to movement is significantly greater than in the opposite direction. In contrast to previous studies, which suggest that brain-wide fluctuation is driven by spontaneous movements (24, 26, 27), our study proposes that the fluctuation, at least in the low-frequency component, is more of an intrinsically organized pattern. Whereas there is no report in vertebrates thus far, a study in *C. elegans* observed a distributed motor command network activated even in the immobilized worm, suggesting brain-wide behavior-related activities independent of re-afferent sensory input (65). If this applies also to the vertebrates, a possible mechanism unifying the seemingly conflicting findings is that the medial brain (the focus of this study) may provide the motivation that guides behaviors (66, 67). This motor guidance is broadcasted to other brain areas and gets manifested in detail by interacting with local neural activities. The movement driven neural activity pattern has been mainly observed in the visual cortex (24); it would be interesting to investigate the temporal coordination between the medial brain and those sensory areas for spontaneous behaviors to understand how these activities are organized. Another reason for the seemingly different conclusion could be attributed to our different data analysis approach. Previous studies adopted multivariate linear models or cross-correlation estimating the variance explained by movements. Our adoption of PDC method allows us to distinguish direct and indirect flows in the frequency domain; this is not possible with correlation methods (50, 68).

Despite the incompleteness of the brain network that could be measured in this study, the general functional division into the external and the internal agrees with modular divisions of other, more exhaustive connectomes. In the rat cerebral cortex, for example, four modules are identified from a graph of directed synaptic connections (15). The four modules are organized into a topologic core-shell structure with two core modules: M1 containing the visual and auditory areas and M2 including mainly the somatic and visceral sensory-motor areas; and two shell modules: M3 including most of the cingulate cortex and hippocampal formation and M4 containing the olfactory system, infralimbic and prelimbic areas, and amygdala. Our study partially overlapped M2-4 under this framework with additional subcortical areas measured. The external state (#1 & #2 in Fig. 4D) was mainly associated with the activation and deactivation of the motor areas (SMA and preSMA), corresponding to activity belonging to the M2 network. These premotor areas are usually linked to motor planning, and for marmosets, the SMA has coevolved with their exceptional volitional vocal control ability compared to other nonhuman primate species (69). The internal state (#3 & #4) was strongly correlated with activity of the ventral subcortical areas including the lateral septum, preoptic area, and hypothalamus. Although these areas are not included in the four modules, the lateral septum, for example, strongly connects with hippocampus (70), which belongs to the core modules M3 and M4. Another area, vmPFC, was also correlated with the internal state (but not statistically greater than the correlation with the external one). This area, more precisely area 25, is a major player in M4 along the middle wall, and connects strongly with the amygdala (71). Thus, one hypothesis yet to be examined is that the internal state may more extensively recruit brain areas within the M4. In the rat connectome study, M2 and M4 exhibit strong reciprocal connections. This may account for the overall highly coupled functional network (Fig. 3E) and the dominating transitions between external and internal brain states.

Brain fluctuation correlated with physiological signals has been found universally (51, 72–75). However, when using the hemodynamic signal as a surrogate for neural activity, such as in functional magnetic resonance imaging (fMRI) and fUS, the cardiovascular and respiratory signals have been treated as confounds and regressed out in fMRI studies (76). In addition, due to signal distortion and CO_2_ concentration change in the brain brought by respiration, it is also a general practice to regress out global fMRI signals approximating the removal of respiration (77). The fUS method does not have the signal distortion issue but could be influenced by CO_2_ concentration. Although we did not explicitly remove potential global influences in our analysis, the PDC estimate automatically treated independent external sources as noise. Thus, the global effect should be discounted at least to the extent of removing the global average. Another technical difference is that the CBV of the arterioles and capillaries measured with fUS (42) are more similar to CBV-fMRI than BOLD-fMRI (44). CBV signals exhibit a shorter onset time and time-to-peak than BOLD signals in marmosets (78) and correlate linearly with neural activity for a wide range of physiological regimes (79–81). The higher sensitivity of the fUS signal may have contributed to our inference of information flow. However, we do realize that the cerebral blood flow is not necessarily driven by local neural activity but may share common input with the electrophysiological activity from, for example, the rostral ventrolateral medulla (82). How faithfully the fUS signals can be translated into neuronal activity at different locations in the brain requires further investigation. Besides the entangled brain-physiological signal owing to the imaging technique, there is also a neurobiological basis for the correlation between brain and physiological states (72). Multiple cortical and subcortical areas participate in autonomic regulations on respiratory, cardiovascular, digestive, and other systems (73, 75, 83). The temporal relationships between brain activity and the physiological signal are frequency dependent. For example, in mice, theta oscillations precede and are Granger-causal for the variation of respiration rate, whereas it is the opposite for the gamma oscillations (84). Our results support that at frequencies <0.5 Hz, the information mainly flows from brain to heart rate, outweighing the opposite direction. However, we did not exclude that heart rate and respiratory sinus arrhythmia can influence CBV at higher frequencies around blood pulsation (~5 Hz) and respiration rate (~1 Hz) in marmosets.

Intrinsic functional architecture revealed in slow spontaneous fluctuations across the brain is not unique to humans (85) and has been observed across various animal species (86–91). With technologies allowing for higher spatial and temporal resolutions, we are now able to better understand the neural and hemodynamic accounts for behavior on a large scale (92). Using fUS, we revealed how heart rate, a measure of the moment-to-moment variation of energy supply to the body, is coordinated with motor activity, which consumes this energy—via a complex brain network. The brain is at the nexus between internal and external environments. Its functional division into external and internal regulations, and the inseparable nature of these two dynamics, support a view of the default mode network as a dynamic sense-making network that actively integrates external and internal life events (93). This coordination may be a consequence of context-dependent energy optimization. Modern robotics design also follows such an architecture with a “task planning” layer for optimizing objectives and a “motor planning and control” layer for taking actions (94). Questions for future studies are how the brain exploits and reconfigures its intrinsic networks in the face of different internal and external challenges in contexts such as social interaction and learning (95, 96), and how life experiences can shape the energy landscape of network dynamics and give rise to the diversity of behavioral phenotypes (93).

## Methods

### Subjects

All experiments were performed in compliance with the Princeton University Institutional Animal Care and Use Committee guidelines. The subjects used in the study were four adult common marmosets (>2 years old, one female and three males) housed at Princeton University. All four subjects participated in the fUS recording; three of them were also used for ECG and video recording. The colony room was maintained at 27°C temperature and 50-60% humidity with a 12L:12D light cycle. Marmosets had *ad libitum* access to water and were fed with a standard diet. All subjects were acclimated to the experimental environment and positioning at least a month before the formal experiments.

### Surgery

For pre-operative procedures, the animal was placed on a warmed blanket with temperature, pulse, respiration and SPO2 being monitored and blood glucose measured. Dexamethasone 1 mg/kg and Baytril 5 mg/kg was administered IM. The animal was induced with alfaxalone 10 mg/kg IM before intubation. The intubated animal was then connected to the anesthesia machine and moved into the OR. The surgical site was clipped, and the remaining hair was removed with Nair. The surgical site was cleaned with betadine and covered with surgical drape (3M). Didocaine was injected into the scalp at multiple sites. An incision (~20 mm, rostral/caudal orientation) was made to expose the skull along the top of the head through the scalp. Tissue was reflected with a retractor to expose the skull. Sterilized miniature titanium screws were inserted into the bone at several locations to anchor the head post and the head plate. A customized head post and a head plate were immobilized on the skull with adhesive cement (C&B Metabond Quick Adhesive Cement System, Parkell). The head plate allowed the ultrasound probe to be aligned with the midline of the brain. Within the head plate, a cranial window of ~8 mm × 16 mm was created. During off-experiment time, a stainless-steel cover was attached to the head plate to protect the head plate and the craniotomy.

### Functional ultrasound (fUS) imaging

Our custom ultrasound probe allowed a wide field of view (20 mm depth, 16 mm width), a temporal resolution of 2 Hz, a spatial resolution of 125 μm in width, 130 μm in depth, and 200-800 μm in thickness depending on the depth (200 μm at 12 mm depth). The probe was connected to an ultrasound scanner (Vantage 128, Verasonics) controlled by a PC. fUS signals were acquired at the midline sagittal plane. The image acquisition method was described previously (44).

### Experimental protocol for fUS recording

Each subject participated in experiments once per day around the same time of the day. The subject was brought to the experimental room from their home cage. The walls of the room (3.2 m × 5.5 m) were covered with sound attenuating foam. The subject was placed in a custom-designed partial restraint device with the head fixed by the head post. The cover was removed to expose the cranial window, which was then flushed with sterile water and covered by sterile ultrasound gel (Sterile Aquasonic 100 Ultrasound Gel). A customized probe holder was then attached to the top of the head plate with screws and the ultrasound probe was placed inside the holder. The head post was released before the experiment to allow head movement. An initial image was acquired to examine the imaging position and quality. Each task-absent trial lasted 10-20 min. After the experiment, the recording surface was cleaned with 0.05% chlorhexidine and sterile water and was closed with the cover. The animal was then returned to their home cage.

### Image processing

We first aligned image frames using a rigid body transformation within each session to eliminate any slow drift of image position within the FOV. To achieve this, we first calculated an averaged image across the session and saved it as a reference. Then we aligned each frame to this reference using elastix software (97). To align all sessions to a common reference, we next consecutively performed rigid body, affine, and B-spline transformations using elastix. We then created a mask for the within brain area shared across all sessions with the sagittal sinus excluded.

To remove timepoints contaminated by motion artifacts, we used the averaged signal of an area outside the brain as control. We set a criterion for a timepoint to be an outlier if the control signal was above 1.5(Q3-Q1)+Q3, where Qi stands for the i^th^ quartile of the control signal. Once the noise was removed, we band-pass filtered the image data between 0.005 Hz and 0.5 Hz and standardized the signal of each pixel. Finally, we spatially smoothed the images using a 2D Gaussian filter with sigma=2 pixels in each dimension.

### Brain parcellation

We first vectorized the image data such that each column was the masked pixels within a frame. To perform group parcellation across all sessions, we calculated principal component analysis (PCA) to reduce the temporal dimensionality (# of columns) to 30 for each session and concatenated all sessions by the reduced dimension (98). Suppose we have the vectorized and centered fUS data for the i-th session 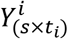, where *s* the number of pixels (the same across sessions) and *t_i_* the number of frames in session *i*. Conducting PCA decomposition, we have 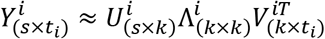, where *k* the number of PCs, *U* the set of spatial eigenvectors and *V* the temporal ones. Projecting *Y^i^* onto the first *k* = 30 temporal eigenvectors, we have reduced data of the i-th session 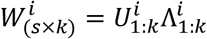. Concatenating *W^i^* of all *N* sessions, we have a group level PCA *W*_(*s×kN*)_ = [*W*^1^,…, *W^N^*]. Brain parcellation was carried out on *W* where the *kN* reduced dimensions were features. Pixels with similar patterns were identified using spectral clustering. Briefly, we calculated the pairwise Spearman’s rank correlation coefficient for the rows of *W*, based on which we established an *s* × *s* similarity matrix *S* with diagonal elements 0 and off-diagonals 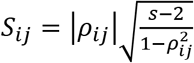, where *ρ_ij_* the Spearman’s correlation between pixel *i* and *j*. We then performed spectral clustering (99) using the similarity matrix for which we calculated the symmetric normalized Laplacian matrix 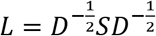, where *D_ii_* = ∑*_j_S_ij_* the degree matrix. We calculated the eigenvectors corresponding to the largest *k* eigenvalues of *L* and performed k-means clustering on the eigenvectors. This yields the spectral clustering results. Similar procedure was carried out for the subject specific parcellation.

### Cardiac data acquisition

To record ECG, we put on an elastic band to the chest of the marmosets with a pair of Ag-AgCl surface electrodes (Grass Technology) sewed on. The data was resampled into 1 kHz and band-pass filtered between 15 and 100 Hz. To identify heartbeats, we calculated sliding correlation coefficients between session-specific templates and the ECG signal, and these locations were marked as one if the correlation coefficient was above a threshold. This binary signal was convolved with a Gaussian window to estimate the momentary heart rate.

### Behavioral videography

In a lighted room, the subject was placed in a customized chair with its front facing a camera (Logitech C920). Videos of the subjects were recorded at 24-Hz framerate and synchronized with fUS acquisition. Essentially, a 1 μs pulse signaling the end of image acquisition (400 ms long per 500 ms cycle) was passed to the interrupt channel of an Arduino board, which converted the signal to a 200 μs TTL pulse to be recorded. The sound was also recorded from a microphone placed next to the subject (Zoom H4n Pro). Audio, TTL pulse, and ECG signal were simultaneously recorded through a data acquisition box (Muscle SpikerBox Pro) at a sample rate of 44.1 kHz. This audio signal was aligned with the audio channel of the video through a brief playback of white noise. The *t*_0_ of ECG and video was aligned to the onset of the first fUS frame’s TTL pulse before downsampling to 2 Hz, where the first sample was kept as the one corresponding to *t*_0_ before the downsampling. We created masks for the subject based on pixel variance contrasting the static background and aligned the masked images across sessions through rigid body transformation and affine transformation using elastix. Video analysis was carried out for the aligned videos. We defined motion intensity (MI) as the absolute value of the total intensity difference between two adjacent frames, scaled by the mean across the session.

To extract different types of motor behaviors, we performed a group independent component analysis (ICA). First, we carried out group PCA by concatenating the first 100 PCs across 35 video sessions. Next, we performed ICA on the group PCs and identified 34 spatial modes. By inspecting the spatial patterns, we categorized the behaviors into six types: blink, mouth movement, ear movement, arm and hand movement, foot and tail movement, and body movement. We calculated the momentary behavioral index as the inner product between each IC and the video frame, then took the maximal value at each time point across the subset of ICs belonging to the same behavioral category. To estimate the fractional time spent on each type of behavior, we calculated the proportion of time during which behavior indices were above a threshold of the session. Time not spent on any movements was determined as being still.

### Coherence

The coherence between motion intensity and heart rate was calculated using the MATLAB function *cmtm*. We concatenated the time series of all 35 sessions to minimize the edge effect. To remove the baseline difference in heart rate by sessions, we used the z-score of heart rate. Statistical significance was determined as being greater than the 95% CI of the null coherence, which was estimated by phase-randomized surrogates of the time series repeated 1000 times. We separated periods of substantial movements and subtle movements using a threshold of MI. If a substantial movement was detected, we also included the timepoints 10 s before and after. The rest was considered to contain only subtle movements. The coherence analyses were repeated for concatenated segments of substantial and subtle movements.

### Partial directed coherence

Partial directed coherence (PDC) infers causal relationships based on the vector autoregressive (VAR) model coefficients in the frequency domain. In the follows, we first introduce the definition of PDC and then modify it to block-PDC (bPDC). Let *x*(*t*) = [*X*_1_(*t*), *x_k_*(*t*)]^T^ be a K-dimension vector time series. In this study, each dimension was the signal of an ROI, the motion intensity, or the heart rate. The VAR model of signal *x* is given by

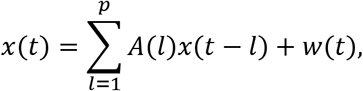

where *w*(*t*) = [*W*_1_(*t*),…, *w_k_*(*t*)]^*T*^ the multivariate zero mean white noise process with covariance matrix *E*[*w*(*t*)*w*(*t*)^T^] = Σ_*w*_, and *A*(*l*) is a *K* × *K* matrix. Here we assume that signals are zero-mean and standardized. To choose the order for the VAR(p) model, we used the Akaike’s information criterion (AIC)

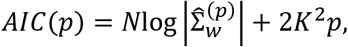

where 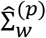 is the estimated covariance matrix with model order *p*, and *N* the number of observations. The maximum likelihood estimator for Σ_*w*_ with order *p* is 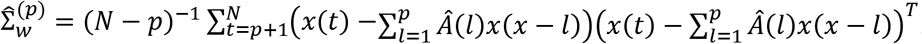. With the optimal *p* selected by AIC, the VAR coefficients *Â*(*l*) were estimated using the Nuttall-Strand method.

Denoting the elements of *A*(*l*) by *a_ij_*(*l*), we define elements of *Ā*(*f*) in the frequency domain

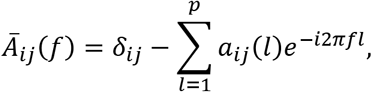

where *f* ∈ [−1/2,1/2). The PDC from *j* to *i* in the frequency domain is then defined as

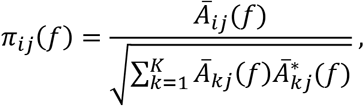

* denotes the complex conjugate. We used the squared PDC measure *π_ij_*((*f*) |, which relates to the mutual information rate via 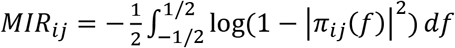. In this work, we used a similar measure as MIR for total information flow between two nodes 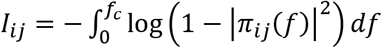, where the cutoff frequency *f_c_* = 0.5 Hz (1/4 unit frequency with a sampling rate of 2 Hz).

To consider the multivariate brain activity as a whole subset of the brain-motor-physiology network, we adopted the bPDC. If *X*(*t*) = [*X*_1_(*t*)^*T*^…,*X_k_*(*t*)^T^]^T^, where *X_k_*(*t*) = [*x_k,1_*,…,*x_k,M_k__*] and 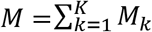 the total dimension of *X*(*t*), is a vector consisting of *K* subsets. Following the VAR representation above, we now divide *A*(*l*) into *K* × *K* blocks *A_ij_*(*l*), each a *M_i_* × *M_j_* dimensional matrix, and hence *Ā_ij_*(*f*) now is defined in the block sense, i.e., the *i* – *j* block. Subsequently, the inverse spectral density matrix of the multivariate time series *X_j_* is

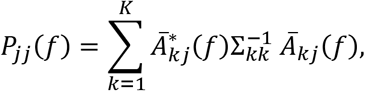

where 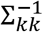 is the *k* – *k* block of the inverse covariance matrix 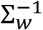. The bPDC from *j* to *i* block is then defined as

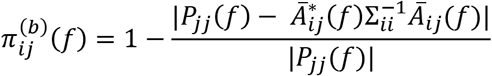

| … | is the determinant. The calculation of the information flow between blocks is the same as the PDC case.

In this study, we concatenated the time-series across all sessions to reduce potential edge effects and increase statistical power. Before the concatenation, we normalized ROI signals and HR, but not MI, within each session. The PDC was calculated using a package available at https://www.lcs.poli.usp.br/~baccala/pdc/. The bPDC expansion is available at https://www.lcs.poli.usp.br/~baccala/pdc/canon/. To test the significance of bPDC values, we phase-randomized the time-series 1000 times and constructed the 95% CI of the null distribution. We set the criteria for statistical significance with Bonferroni correction for multiple comparisons *α* = 0.05/6, as there were 6 directional relationships to examine. To test the difference between *I*_*i*_1_*j*_1__ and *I*_*i*_2_*j*_2__, we bootstrapped the data by resampling sessions with replacement and calculated bPDC for each resampled time series. This procedure was repeated 1000 times to estimate the distribution of information flow in each direction. We then compared each pair of directions using paired z-test with a Bonferroni corrected criterion *α* = 0.05/15.

We defined information flow above the 75th-percentile within each category as strongly influencing directions. The categories were 1) *I_ij_* for *i,j* ∈ *ROIs*, 2) *I_ij_* for *i* ∈ *ROIs*, *j* = *MI* and *i* = *MI*, *j* ∈ *ROIs*, and 3) *I_ij_* for *i* ∈ *ROIs*, *j* = *HR* and *i* = *HR*, *j* ∈ *ROIs*.

### Brain image registration to atlas

We adoped a two-step registration using the volumetric brain vasculature images as a nexus linking the fUS images to a standard brain atlas. More details of this procedure (applied to a rat brain) has been described in ref. (100), here we briefly introduce the steps and the adaptation for marmoset brains.

#### Perfusion

After the endpoint of each subject, the brain was perfused with 50 ml fluorescence-tagged albumin hydrogel. The hydrogel was made of 5 mg AF647-albumin (A34785 Invitrogen) mixed in 50 ml 2% (w/v) gelatin (G6144 Sigma-Aldrich) in PBS. The brain was extracted and further submerged in 4% PFA at 4°C overnight, and then was processed using iDISCO clearing protocol (101).

#### Light-sheet microscopy

The brain was cut along the sagittal direction into six pieces of equal thickness (~4 mm) with the cerebellum excluded. Each chunk was imaged with the medial side facing up under a light-sheet microscope (LaVision Biotech) using a 1.3x objective. We used the 488 nm channel for autofluorescence imaging and the 647 nm for vasculature imaging. The sample was scanned with a 1.3x objective, which yielded a 5 um x 5 um pixel size and with a z-step of 5 um.

#### Registration

The registration procedure took two steps: one registered the autofluorescence channel of light-sheet images to a standard brain atlas (Step 1), and one registered a reference fUS image to a plane of vasculature light-sheet image from the same subject (Step 2).

##### Step 1: light-sheet to the brain atlas

The light-sheet image stacks were first down-sampled to 50 um isotropic. Using the autofluorescence images, the surfaces between two adjacent brain pieces were extracted and matched using iterative closest point algorithm (MATLAB routine *icp*). The chunks were translated and rotated numerically to be coarsely assembled. We then registered this assembly to a brain altas (102) by applying rigid and affine transformations using Elastix (97). After this coarse registration, we refined the alignment to the atlas for each piece iteratively by applying rigid, affine and B-spline transformations. The same transformations were applied to the vasculature channel, thereby registering the brain vasculature to the atlas.

##### Step 2: fUS to light-sheet

To localize the fUS position in the brain, we created a subject-specific local 2D vascular atlas by taking the max-intensity projection over 300um thickness along the midline of the registered vasculature volume. The fUS image of the same subject was aligned with this vasculature reference using a landmark-based 2D registration procedure (ImageJ plugin bUnwarpJ(103)). The fUS images were hence registered to the standard atlas.

### Directed graph clustering

To detect clusters (or communities) within a directed graph, we adopted the weighted cuts algorithm for spectral clustering (104). Here the similarity matrix *S* is the asymmetric directed information flow with zero diagonal between each pair of nodes in the network, i.e., the ROIs. Essentially, we normalized the Laplacian matrix with node weights *T*; here we used *T_ii_* = *D_ii_* = ∑_*j*_ *S_ij_*, which is simply the out-degree. We next found the eigenvectors corresponding to the *k* smallest eigenvalues of 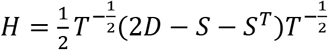, and then performed the k-means clustering on the eigenvectors. The clustering result identifies the nodes belonging to the same cluster. The MATLAB package can be found at https://sites.stat.washington.edu/mmp/software.html.

### Brain state clustering

We followed (60) to decompose imaging data into a spectral-spatial space. Using the group-PCA *W* calculated above, we further extracted 25 group independent components (group-ICA) representing the spatial distribution of potential source signals. We filtered the brain images into spectral bands using maximal overlap discrete wavelet packet transformation with a Daubechies wavelet of 2 vanishing moments. Then we multiplied the ICs and each of the frequency bands of the brain image 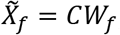, where *C* the ICs of size *n_ic_* × *n_pixel_* and *W_f_* the wavelet packet transform of frequency *f* with size of *n_pixel_* × *n_time_*. Thus, the dimension of a time point was reduced to *n_ic_* × *n_freq_* = 400. For the multitaper approach, we decompose each pixel into time and frequency by calculating multitaper spectrum over a moving window. The spectrogram was estimated using the *chronux* MATLAB package (105, 106) function *mtspecgramc*, with TW=0.5, K = 1, and 4.5 s window size.

One problem with group-level clustering is that variations created by experiments due to slight differences in, e.g., probe positioning and signal quality and the difference in subjects’ brain morphology may become dominant and drive the clustering results. To deal with this, we minimized the dependence on experimental sessions using an expectation-maximization (EM) procedure with penalization on session dependence in the objective function of K-means clustering. Then, we transformed the data iteratively to a common space across sessions. This algorithm was based on the Harmony method developed for high-dimensional biological data like RNA-seq (107). In this study, we modified this algorithm to deal with time series with the temporal structure maintained during the optimization by applying ridge regularization (the elaborated method will appear in a separate article). The transformed data yielded clustering with session-dependence largely reduced (Fig. S10). Data used for the correction were the dimensionality reduced data described above. We further performed k-means clustering with correlation measure for distance to identify brain state for each time point.

### State score

To evaluate to what extend the ROI activity was attributed to changes in brain state, we estimated the percentage variance of ROI activity explained by the occurrence of each brain state using cross-validation. We randomly stratified the data into ten folds, trained linear models of *y_i_* = *α_ij_* + *β_ij_X_j_* + *ϵ_i_* where *y_i_* the activity of the ith ROI, *X_j_* the one-hot encoding of brain state *j*, and *ϵ_i_* a random noise, using the nine folds of the data, and calculated test variance explained as *R*^2^ = 1 – *RSS/TSS* using the remaining fold. We estimated the mean and standard deviation of variance explained across the 10-time cross-validations for each ROI and each brain state. Assuming the variances explained were normally distributed, we tested whether the variance explained was equal to zero. The null hypothesis would be rejected if the mean variance explained was greater than 1.96 times the estimated standard deviation. Our results showed that states #1 and #4 co-varied and #2 and #3 co-varied. We thus defined a state score as 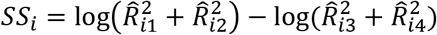, where 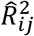 is the variance explained for ROI #i by the j-th state. Thus, if a signal were more strongly correlated with #1 and #2 than #3 and #4, the state score would be positive, and vice versa. To estimate the variance of the state scores, we applied chain rule 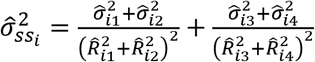.

### State correlation with control preference

We defined the control preference as the difference between the information flow from ROI to movement and from ROI to heart rate: *CP_i_* = *I*_*i*→*move*_ – *I*_*i*→*HR*_. To test the dependence of control preference on brain state, using each ROI as a sample, we performed a weighted linear regression with the weight as 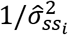. (a similar result was obtained with ordinary least squares). The criterion for statistical significance was set at *α* = 0.05.

### Power spectral density

To estimate the temporal profile of the alternation between the two groups of states, we converted the sequence of brain states into a binary sequence with #1 and #2 labeled 1 and #3 and #4 labeled 0. We used the *chronux* routine *mtspectrumc_unequal_length_trials* to estimate the power spectral density (PSD) and the 95% confidence interval of the PSD with taper TW=10.5 and K=20.

## Acknowledgments

We thank Uri Hasson and Samuel Nastase for comments on the manuscript. This work was supported by an NIH-NINDS grant to A.A.G. (R01NS054898).

## Author Contributions

Conceptualization: YSZ, AAG

Methodology: YSZ, DYT

Funding acquisition: AAG

Data collection: DYT, AEH, YSZ, DAL

Data analysis: YSZ

Writing: YSZ, AAG

## Data and material availability

Data is available from Dryad Digital Repository doi:10.5061/dryad.jwstqjq9x

## Supplementary figures

**Fig. S1.**
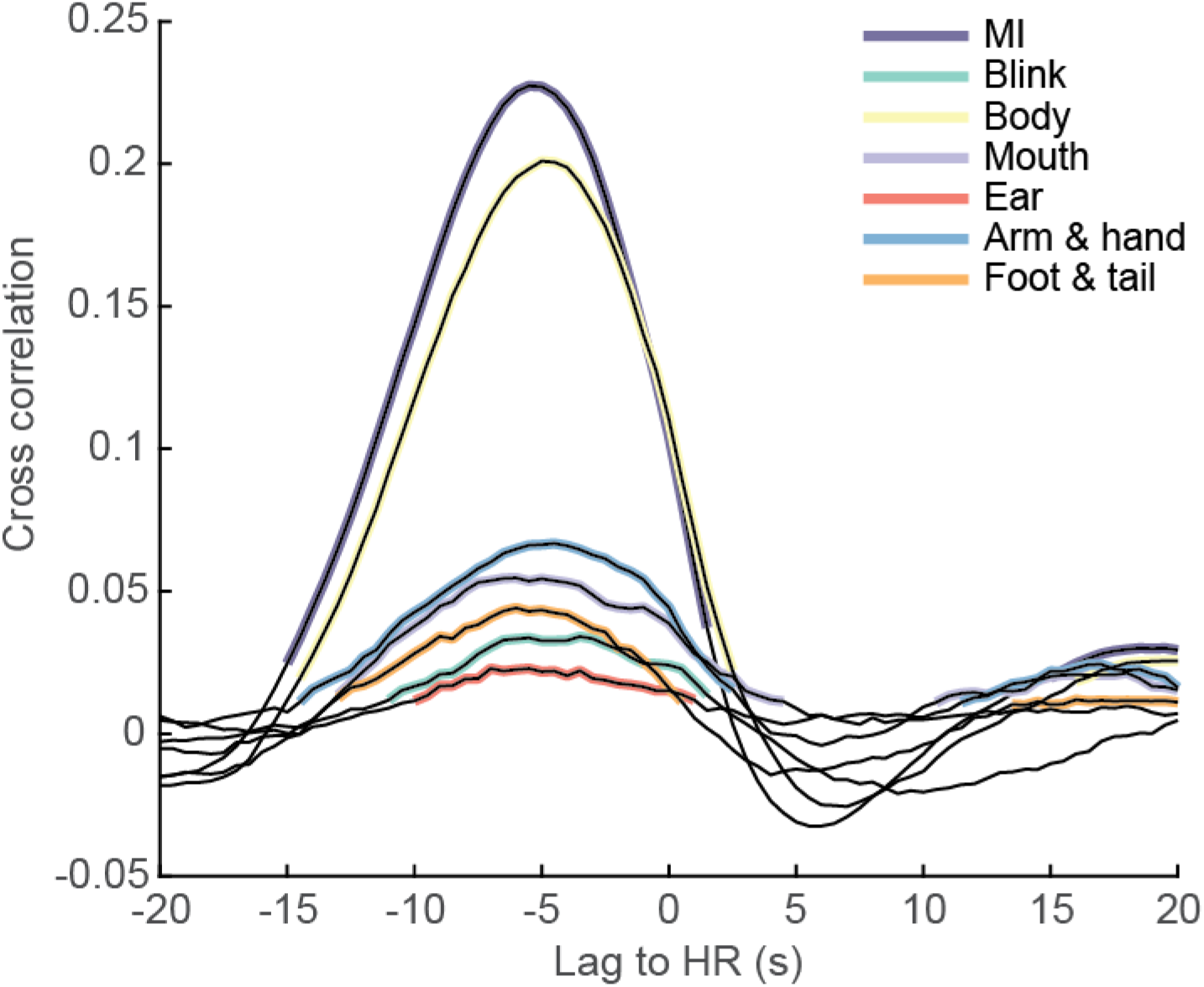
Cross correlation between HR and different types of movements and MI. Highlighted segments are significant against the 95% CI of phase randomized data. Positive lag is HR leading and vice versa. The time lags of different movements and MI are consistent.

**Fig. S2.**
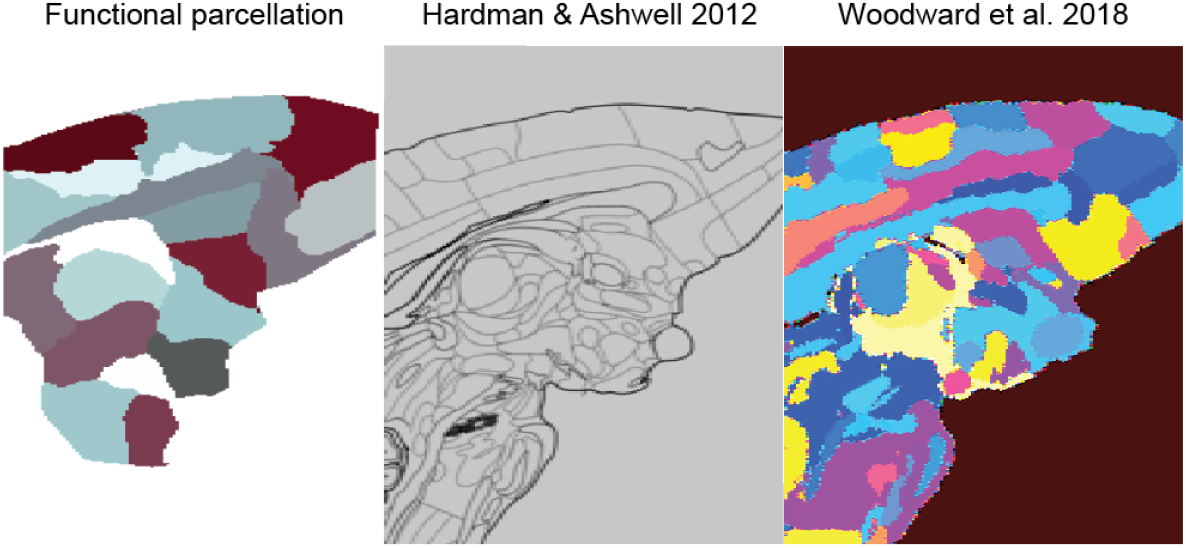
Functional parcellation compared with two marmoset brain atlases.

**Fig. S3.**
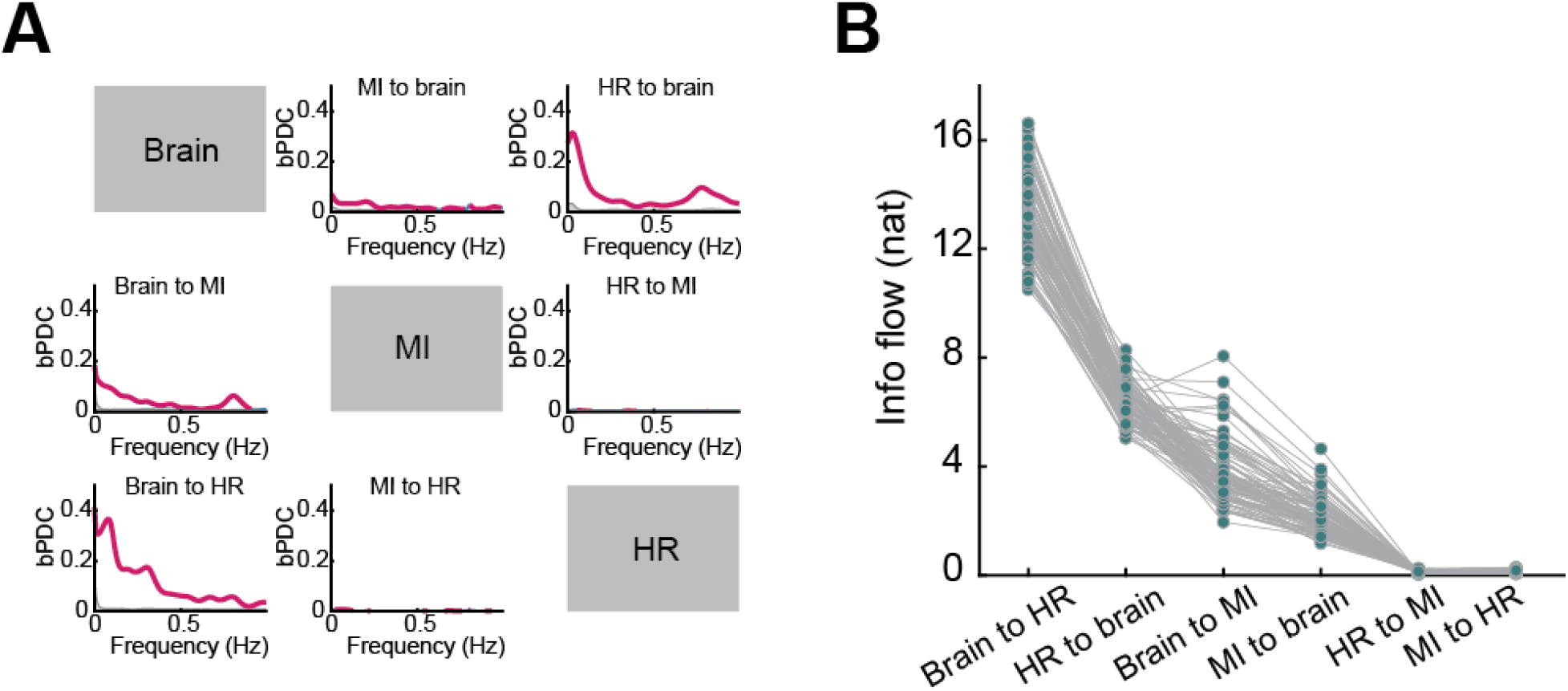
bPDC analysis of un-filtered data. (A) bPDC as functions of frequency for each pair of time series. (B) Bootstrapped distribution for information flow sorted in descending order. Note that the information flow from brain to heart rate (HR) is significantly greater than HR to brain and from brain to movement (MI) is greater than MI to brain. p<0.05, Bonferroni corrected.

**Fig. S4.**
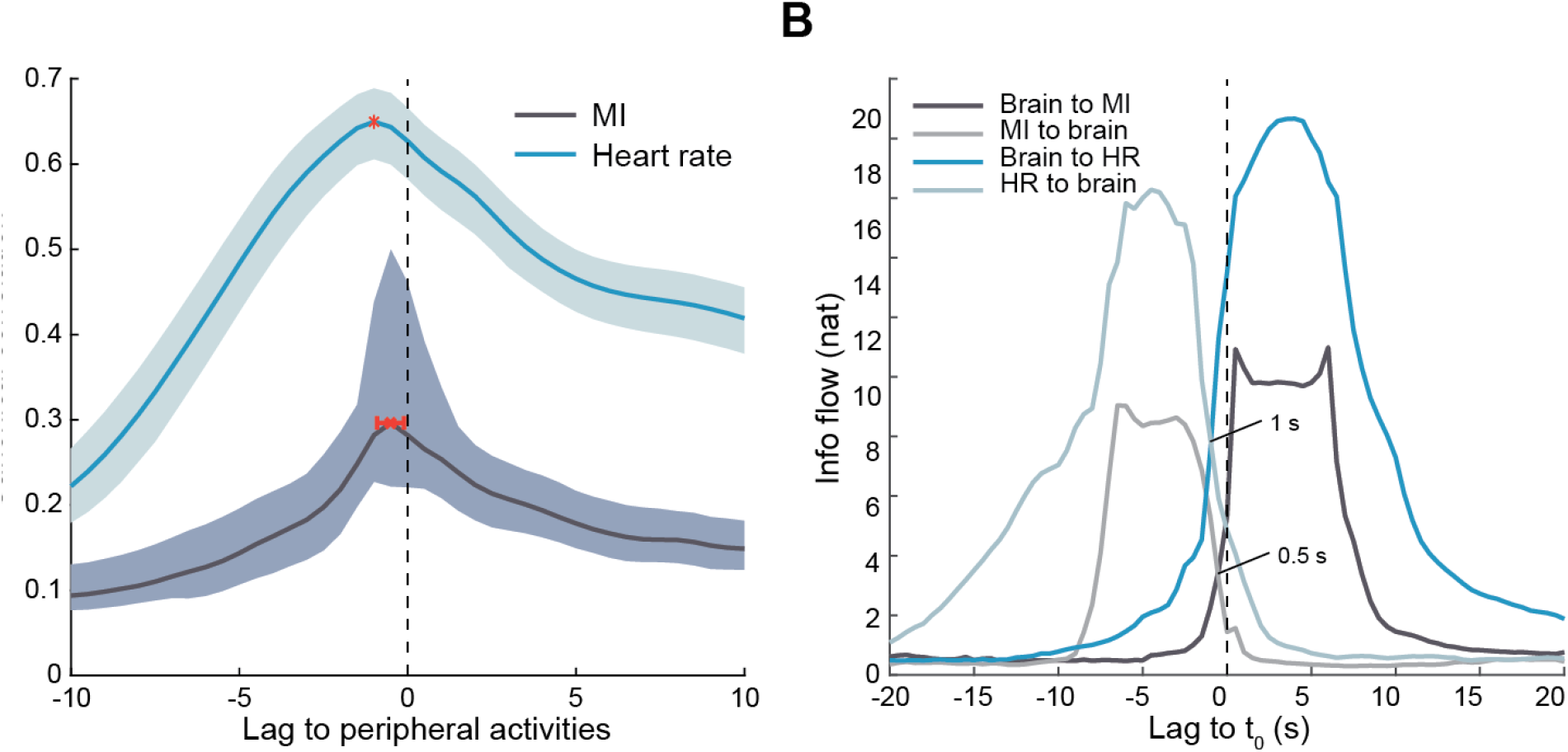
Delay test. (A) Canonical correlations between ROI activities and movement and between ROI activities and heart rate. Shaded areas are 95% CI. Red bars label the peak position. For movement, the lag of brain is −0.51±0.40 s, and for heart rate it is −1.0±0.0 s (mean +\− s.d.). A negative lag means that brain is leading. (B) Information flow with shifted fUS signal to different time delays. Rightward is fUS more ahead. The causal relationship flips when two opposite information flow curves intersect.

**Fig. S5.**
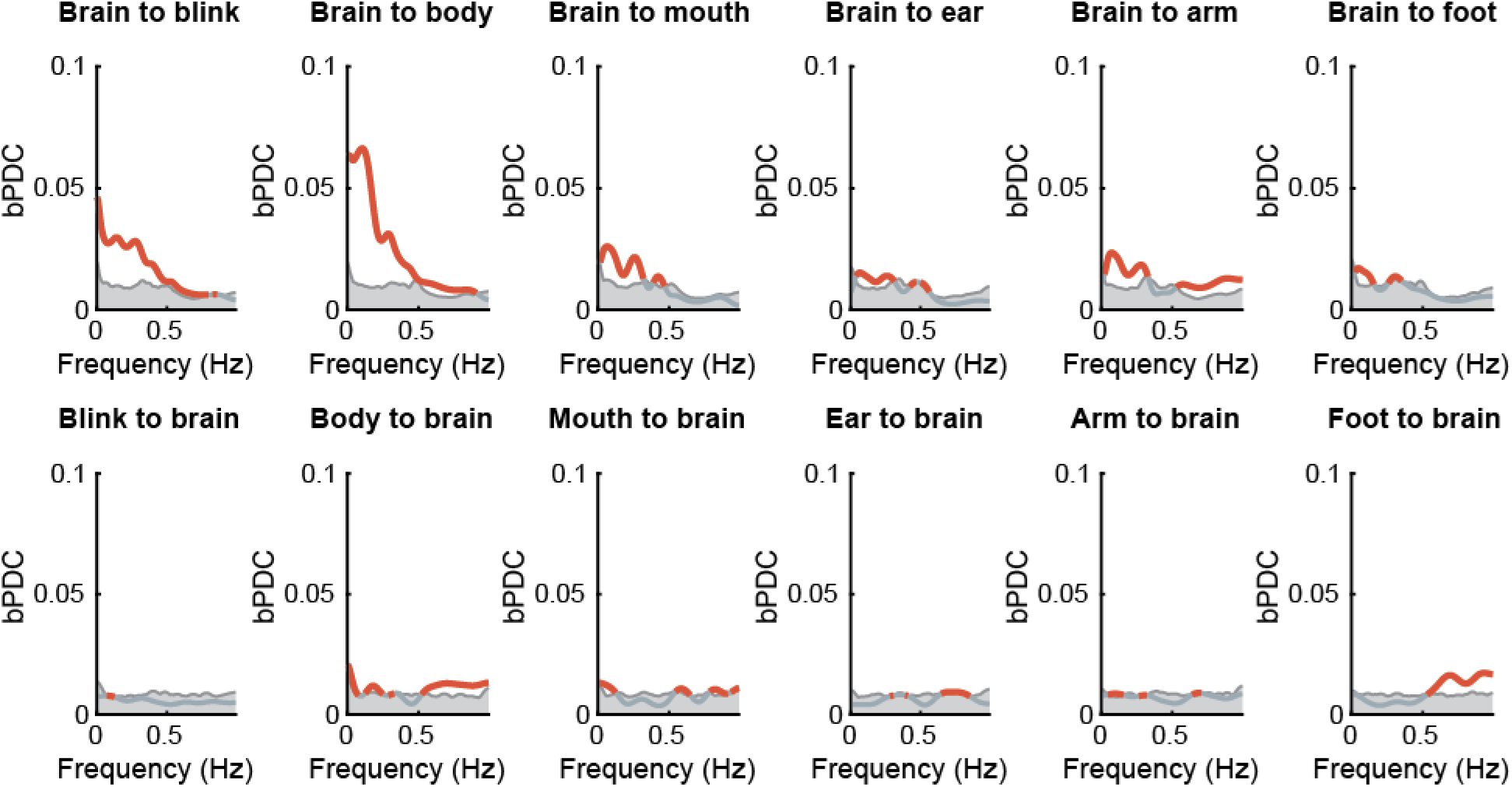
bPDC between ROIs and each type of movement. The bPDC was among three subsets: ROIs, types of movements, HR. Red highlights values significantly higher than the 95% CI of phase-randomized surrogates.

**Fig. S6.**
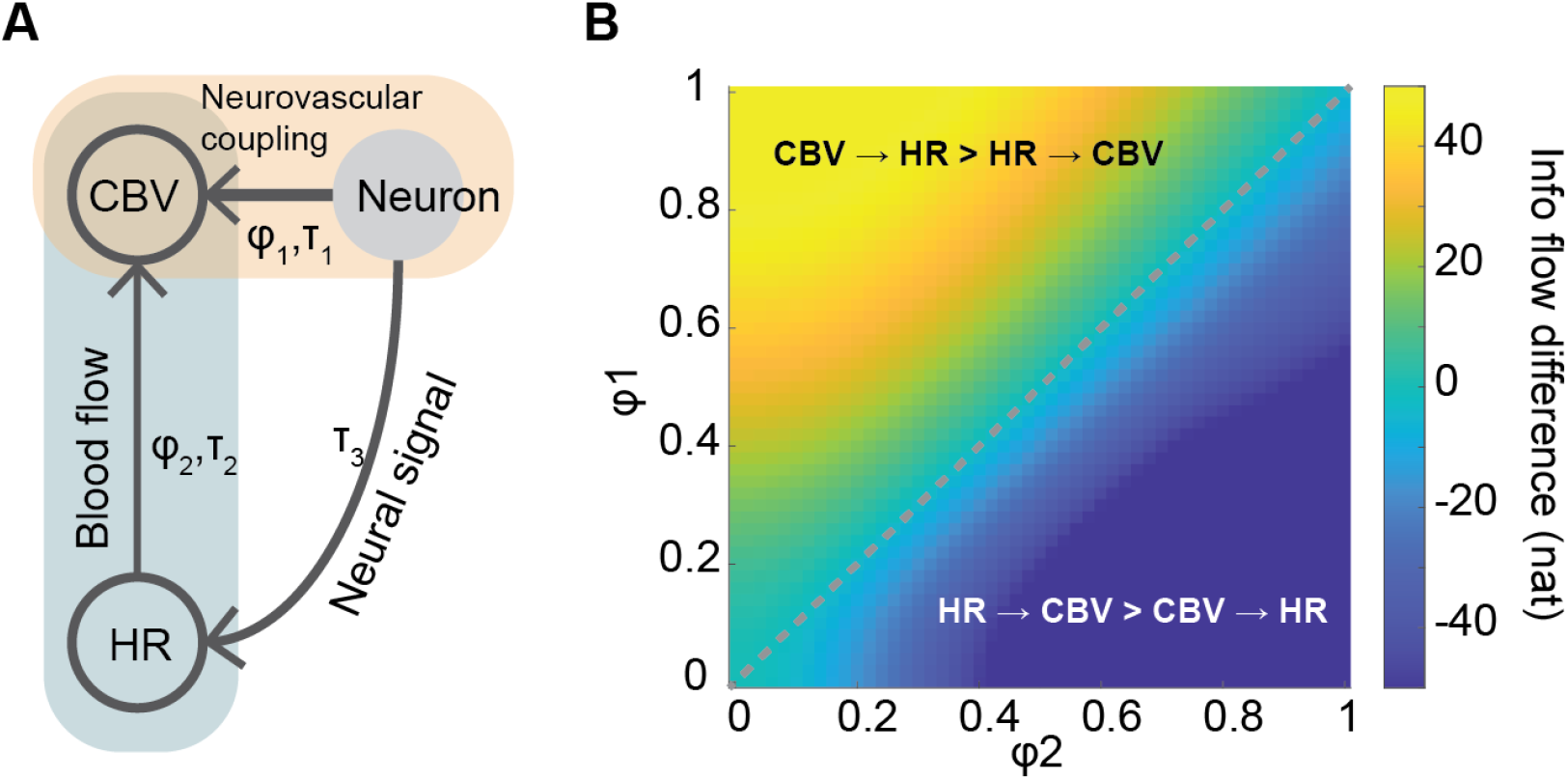
A possible mechanism for information flow from CBV to heart rate (HR). (A) A diagram for CBV, neural activity, and HR relationships. CBV is changed by local neuronal activity via neurovascular coupling with a connection strength *ϕ*_1_. HR influences CBV via brain vasculature with a strength *ϕ*_2_. Neural activity affects HR via a neural signaling pathway. The delay *τ*_2_ > *τ*_1_ and *τ*_3_ > *τ*_1_. Denoting the Neuron, HR, and CBV activities *x*_1_, *x*_2_, and *x*_3_, the model for the simulation is

*x*_1_(*t*) = 0.9901*x*_1_(*t* – 1) – 0.2475*x*_1_(*t* – 2) + *ϵ*_1_,
*x*_2_(*t*) = 0. 25*x*_2_(*t* – 1) + 0.75*x*_1_(*t* – 3) + *ϵ*_2_,
*x*_3_(*t*) = *ϕ*_1_*x*_1_(*t* – 1) + *ϕ*_2_*x*_2_(*t* – 5) + *ϵ*_3_.

CBV and HR are the observations and Neuron is Hidden. (B) The information flow difference between HR and CBV in the space of *ϕ*_1_ and *ϕ*_2_. The upper left triangle corresponds to the regime where we can observe greater information flow from CBV to HR.

**Fig. S7.**
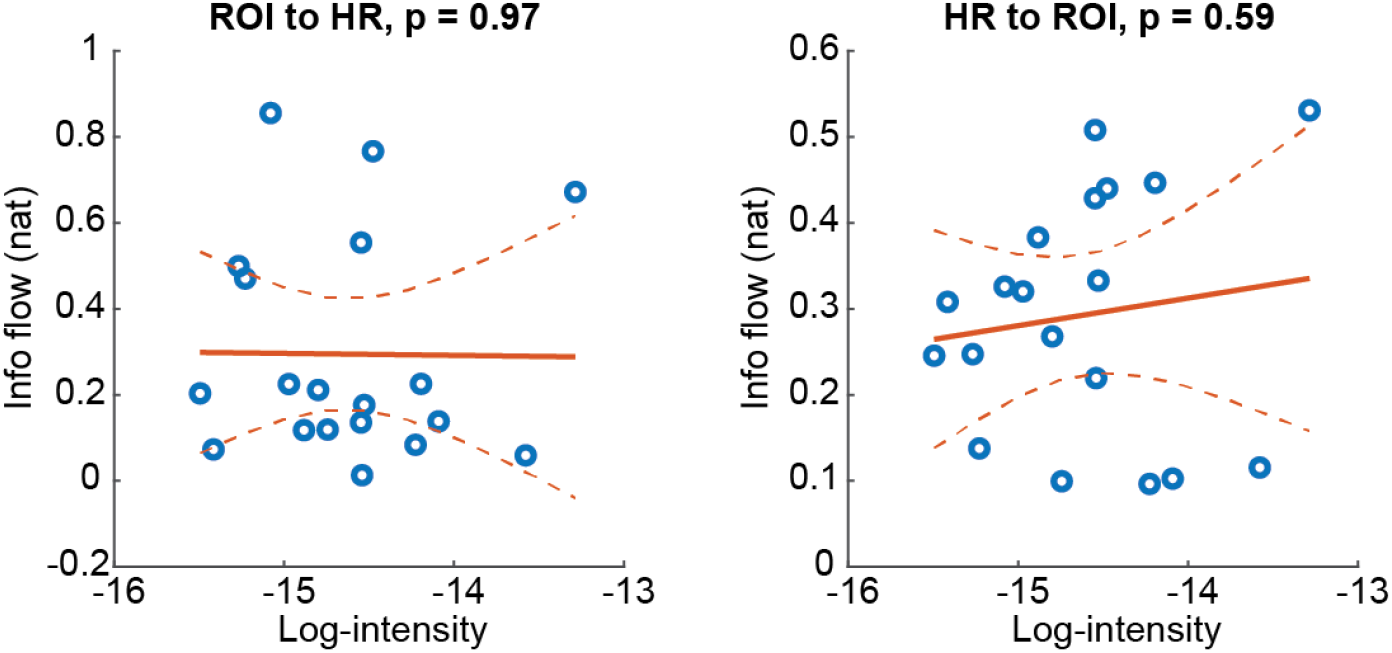
Information flow between ROI and heart rate (HR) vs. image intensity (in logarithmic scale). The fUS intensity is positively correlated with CBV.

**Fig. S8.**
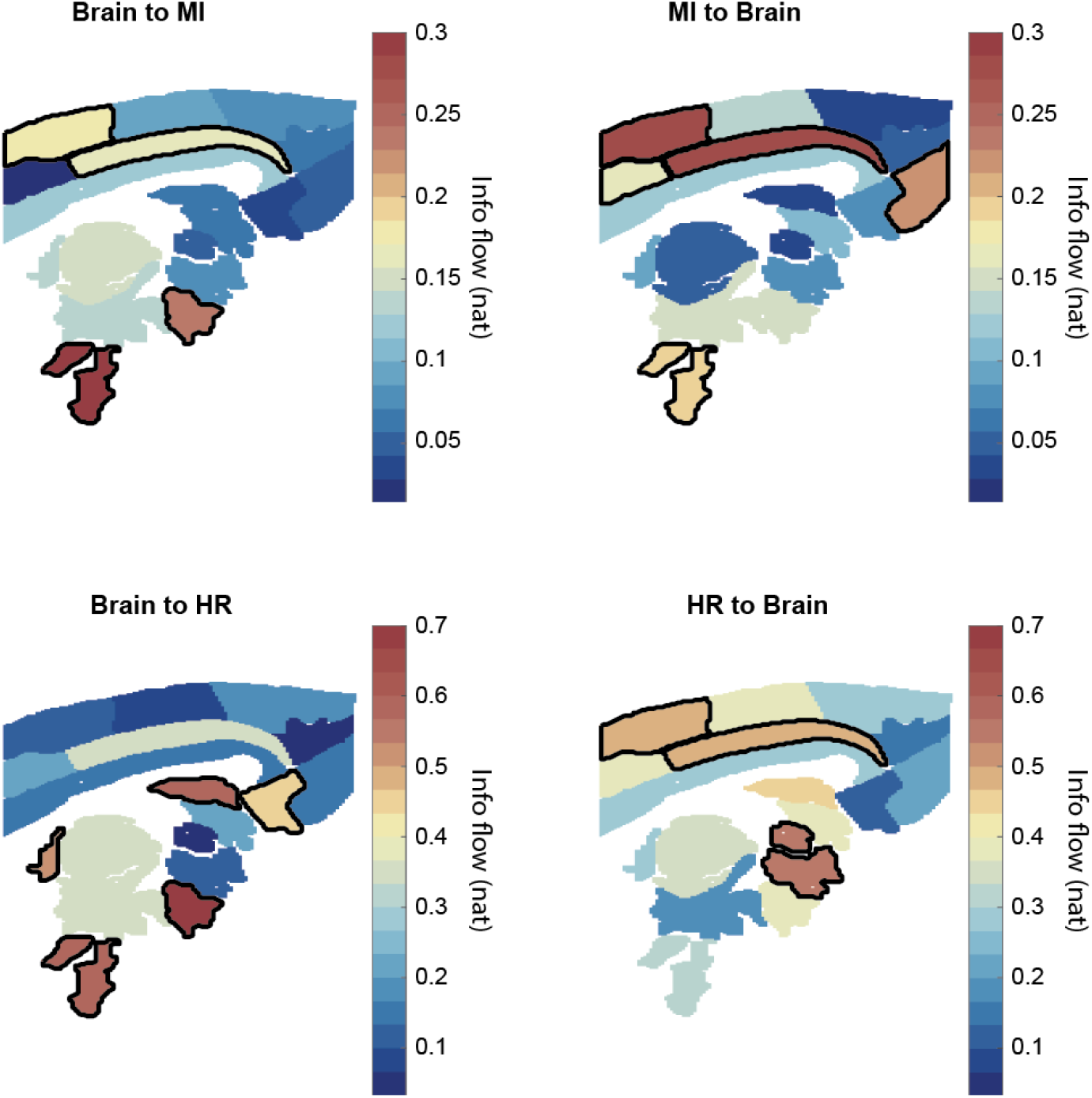
Information flow between individual ROIs and movement (top) and between ROIs and heart rate (bottom). Highlighted areas with bold boundaries are above 75% of the values. The ROI segmentation is based on ref. (48). Note that with this brain parcellation method, the RF and VMH were involved in both movement and heart rate regulation.

**Fig. S9.**
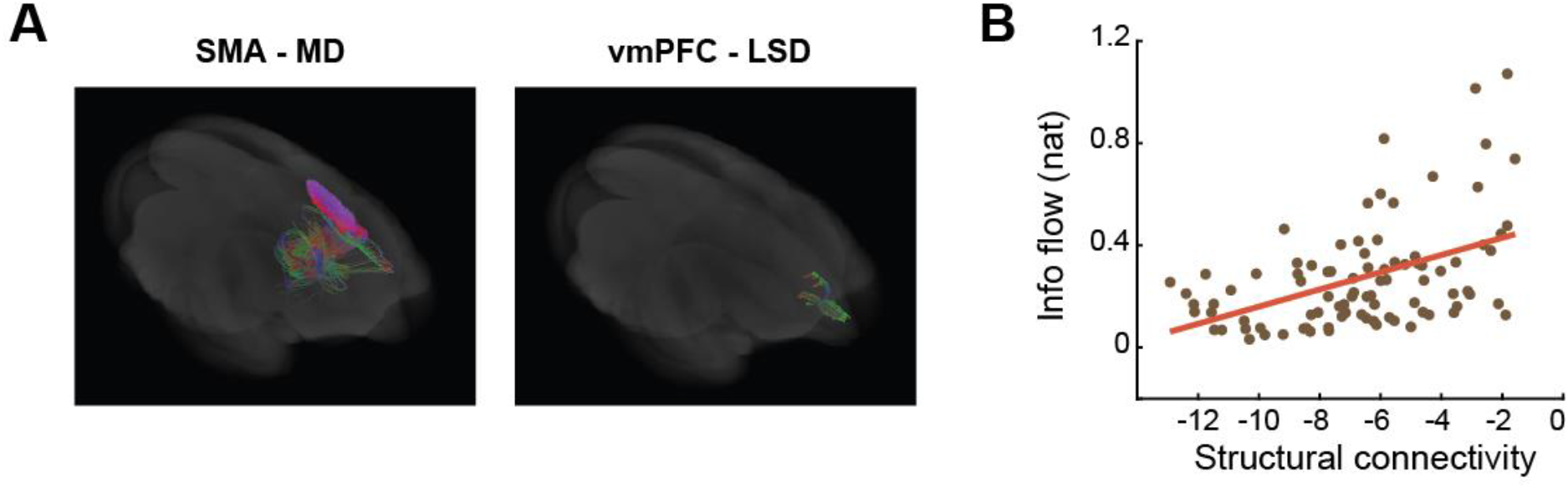
Functional connectivity correlates with structural connectivity. (A) Examples of fiber tractography. (B) Information flow (symmetrized) vs. log-probabilistic tractography.

**Fig. S10.**
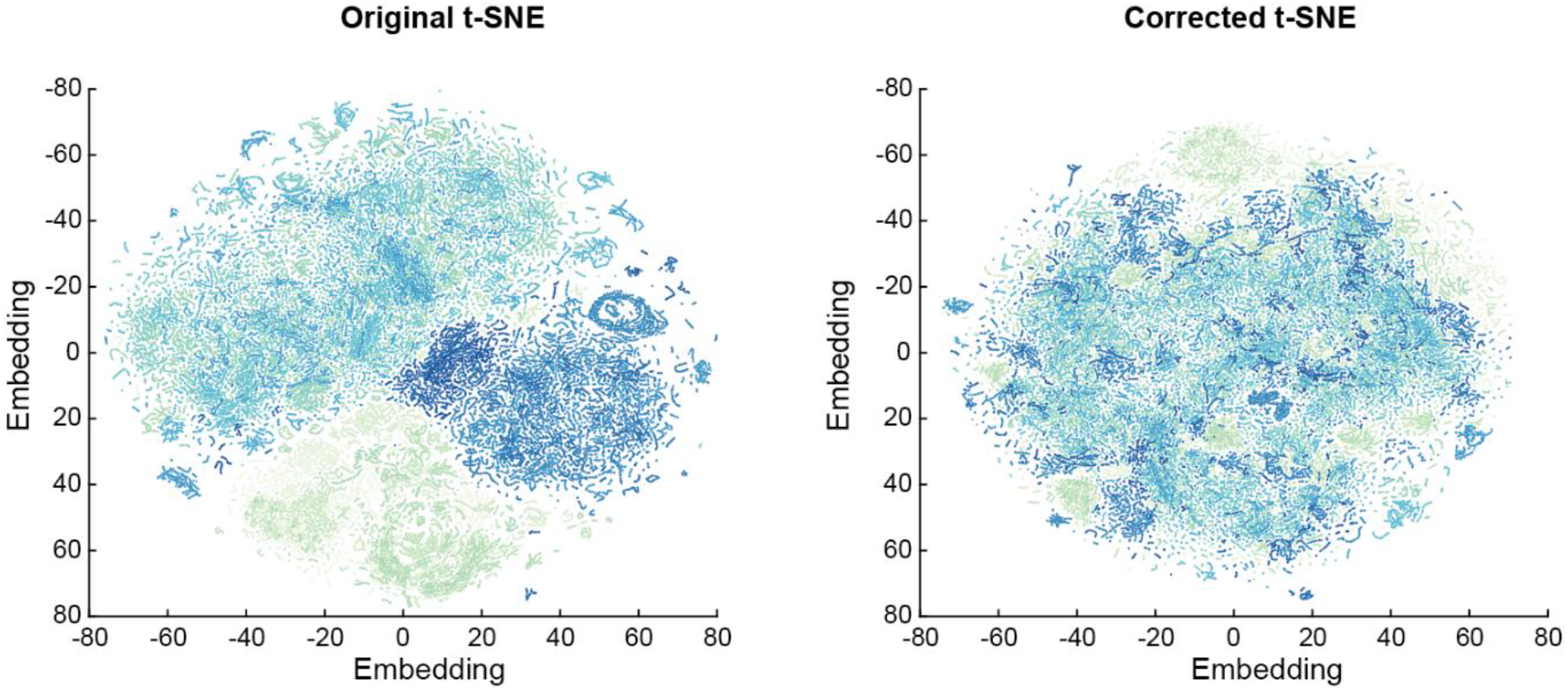
Global brain state visualization using t-SNE before and after correction for session dependence. Color represents session number.

**Fig. S11.**
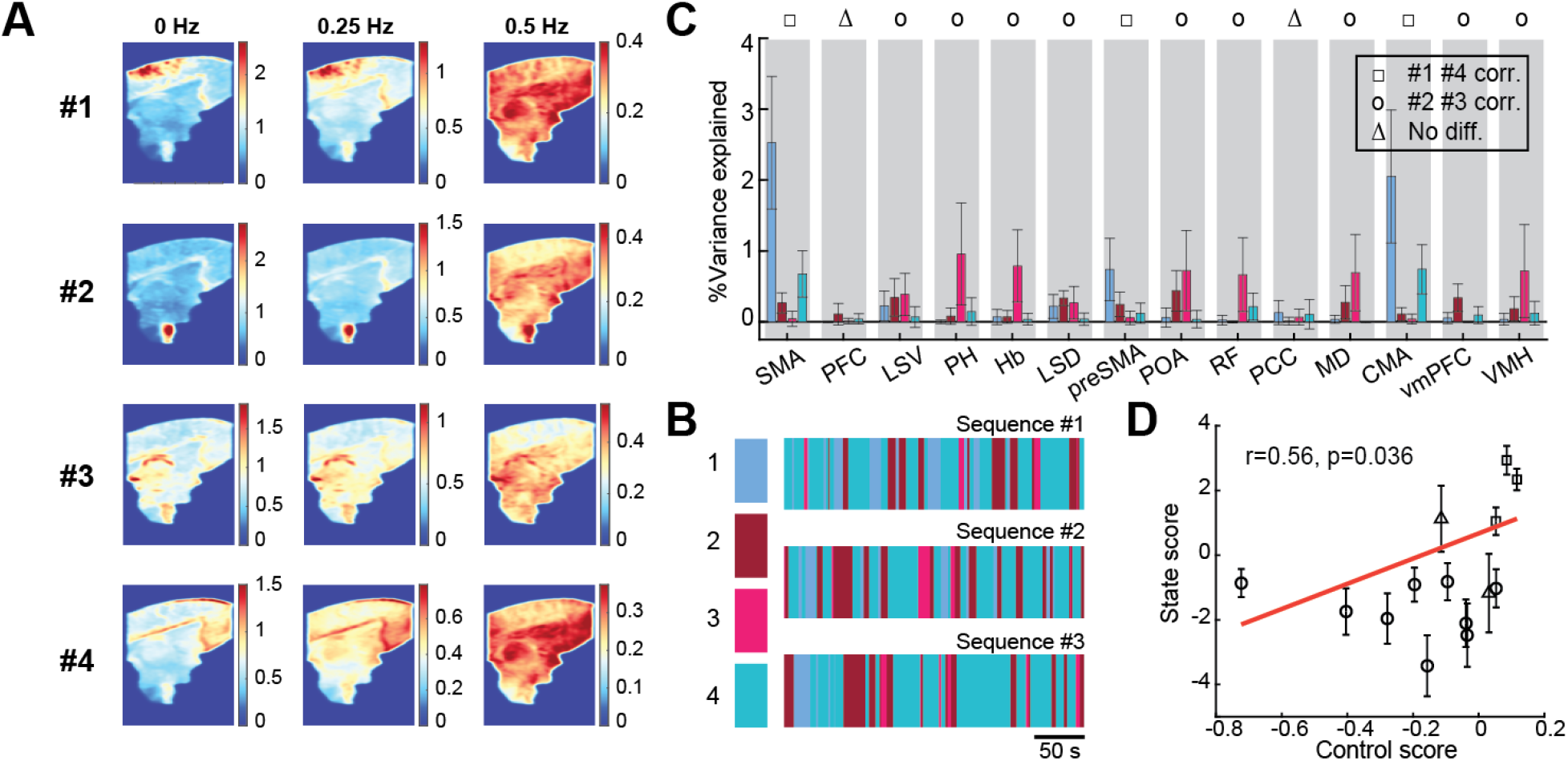
Brain state clustering by ICA-multitaper method. (A) Averaged power spectral density image per cluster estimated by multitaper method. (B) Exemplar state sequences. (C) ROI activity variance explained by each brain state. (D) Correlation between state score vs. control score.

## Supplementary figures

### Proof of PDC’s invariance to causal filtering

Given the Fourier transform of a multivariate time-series *X*(*f*), applying any causal filtering *G*(*f*) gives filtered signals *Y*(*f*) = *G*(*f*)*X*(*f*). In the frequency domain, *X*(*f*) can be written in a moving average representation *X*(*f*) = *H*(*f*)*w*(*f*), where *H*(*f*) stands for the moving average coefficient and *w*(*f*) is the Fourier transform of the zero-mean innovation process. Hence we have *Y*(*f*) = *G*(*f*)*H*(*f*)*w*(*f*). Assuming *H* and *G* are invertible, we can further write *H*^-1^(*f*)*G*^-1^(*f*)*Y*(*f*) = *w*(*f*).

By definition, the PDC from *j* → *i* is 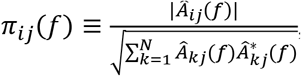, where the matrix *Â*(*f*) = *I* – *A*(*f*) and *A*(*f*) is the autoregression coefficient and *N* the total number of signals. It is known that *Â*(*f*) = *H*^-1^(*f*) and hence we have *Â*(*f*)*G*^-1^(*f*)*Y*(*f*) = *w*(*f*). As *G* is diagonal (otherwise inter-dependence is artificially introduced), element-wise we have 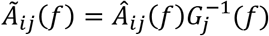. The new PDC of *Y* is then 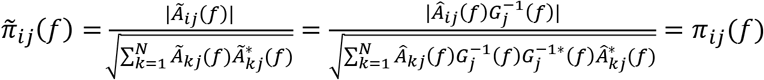. Q.E.D.

